# RiboRep: Replicate-Aware Cross-Modal Transformers for Codon-Resolved Ribosome Density Prediction

**DOI:** 10.64898/2026.08.03.742524

**Authors:** Ai-Te Kuo, Zongliang Yue, Wei-Shinn Ku, Hao Chen

**Affiliations:** Auburn University, Auburn, Alabama, USA

## Abstract

Ribosome profiling enables genome-wide measurement of translation at nucleotide resolution and provides a dynamic view of cellular protein synthesis under diverse biological conditions. Existing computational approaches primarily operate on codon-level representations, potentially losing fine-grained translational signals critical for modeling context-dependent cellular responses. Such predictive translational modeling is increasingly important for emerging biological digital twins, where accurate simulation of molecular-state dynamics is required to characterize cellular adaptation, perturbation response, and phenotype progression. We present RiboRep, a replicate-aware cross-modal transformer for codon-resolved ribosome density prediction. RiboRep jointly models nucleotide-resolution RNA sequences and reference ribosome occupancy signals using dual-stream convolutional encoders, RoPE-based self-attention, asymmetric cross-attention, and replicate-aware conditioning tokens. By explicitly modeling replicate-specific variation and integrating sequence context with experimentally observed translational activity, RiboRep provides a framework for reconstructing and simulating translational states across biological conditions. Across bacterial, yeast, and plant ribosome profiling datasets, RiboRep achieves competitive or improved performance compared with existing baselines, with particularly strong gains on replicate-rich plant datasets. Ablation studies further demonstrate the importance of local codon-aware feature extraction, replicate-aware conditioning, and gated readout. Beyond predictive performance, the proposed framework establishes a foundation for translation-aware molecular digital twins capable of modeling ribosome occupancy landscapes, perturbation-induced translational responses, and condition-specific regulatory programs at codon resolution. ^1^

## 1 Introduction

Ribosome profiling has emerged as a powerful genome-scale technology for quantifying translation by sequencing ribosome-protected mRNA fragments, enabling near-nucleotide-resolution measurement of ribosome occupancy across coding regions [1, 2]. In the context of molecular and cellular digital twins, ribosome profiling provides a critical dynamic layer between transcript abundance and protein-level phenotype, because it captures how cells allocate translational capacity across genes, codons, and regulatory contexts. Rather than treating gene expression as a static transcriptomic readout, Ribo-seq enables the construction of translation-aware computational models that can simulate ribosome movement, elongation heterogeneity, pausing, and condition-specific translational responses. Ribosomal occupancy patterns encode regulatory information that is directly relevant to digital twin modeling of cellular state transitions. Local changes in ribosome density can reflect codon usage, mRNA sequence context, ribosome pausing, ribosome collision, tRNA availability, nascent peptide effects, and stress- or perturbation-induced remodeling of protein synthesis [3]. These features provide mechanistic signals for predicting how molecular perturbations propagate from RNA sequence and translational regulation to protein production and downstream phenotypic outcomes.

Accurate prediction of ribosome density from mRNA sequence and Ribo-seq data is therefore an enabling component for translation-aware biological digital twins. Such models can serve as in silico systems for reconstructing baseline translational states, simulating perturbation-specific ribosome occupancy landscapes, identifying cis-regulatory elements that control translational output, and prioritizing disease- or stress-associated translational mechanisms. Recent advances in deep learning have enabled prediction of ribosome occupancy from sequence and ribosome profiling. Recent deep-learning approaches, including RiboMIMO [4], Riboexp [5], and Riboformer [6]have been developed for ribosome-density modeling. These approaches primarily operate at codon-level resolution, aggregating nucleotide signals into triplet windows. We hypothesize that maintaining nucleotide-resolution inputs before codon-level aggregation can capture finer translational dynamics and enable discovery of sequence determinants associated with reading frame, codon periodicity, and local sequence context.

In this work, we introduce RiboRep, a replicate-aware transformer framework for ribosome density prediction from RNA sequence context and reference ribosome-density profiles. RiboRep jointly models nucleotide sequence, reference occupancy signals, and replicate identity to predict target ribosome density at codon resolution. Unlike existing approaches [4–6], which operate on codon-resolution representations, RiboRep models the nucleotide-resolution sequence and density input prior to codon-aware aggregation. Our contributions are threefold. First, we preserve nucleotide-resolution sequence and density inputs and use branch-specific convolutional encoders to extract codon-aware local features before global attention. This design preserves fine-grained nucleotide information while producing biologically meaningful codon-aware representations. In this paper, we used a kernel size of 3 to reflect codon-scale structure. Different convolutional kernel sizes may also enable future exploration of alternative local sequence patterns. Second, we introduce a replicate-aware dual-stream sequence-density architecture with asymmetric cross-modal attention. Conditioned on the target sequence representation, the model retrieves informative evidence from the reference-density stream through sequence-density cross-attention, enabling adaptive integration between RNA sequence context and experimentally observed ribosome occupancy patterns. Such conditioning mechanisms have become widely adopted in various domains for context-dependent representation learning [7, 8]. Third, we introduce a replicate-aware conditioning special token [9, 10] in the sequence to allow the model to account for replicate-specific biological and technical variation. Inspired by [5], we adopt a gated readout mechanism to adaptively aggregate position-wise representations for final prediction. Extensive experiments across bacterial, yeast, and plant datasets demonstrate that RiboRep achieves competitive or improved performance compared with existing approaches.

## 2 Method

### 2.1 Problem Formulation

Let X = {*A, C, G, U*} denote the four base types of RNA nucleotides, with optional extension to include ambiguous nucleotide symbols. An mRNA sequence of length *L* is represented as *X* = (*x*_1_, *x*_2_, …, *x*_*L*_), where *x*_*i*_ ∈ X. Let *D* = (*d*_1_, *d*_2_, …, *d*_*L*_) denote the corresponding ribosome footprint density profile at nucleotide resolution, where *d*_*i*_ ∈ ℝ_≥0_ represents the P-site-assigned ribosome occupancy at position *i*.

Given a codon-centered window x_*i*_ ∈ X^*w*^, the reference density profile d_*i*_ is obtained from one replicate within a replicate group and used as input. The target *y*_*i*_ denotes the codon-level value at position *i*, computed from another replicate from a separate replicate group. Our goal is to learn model parameters *θ* by minimizing the following loss:

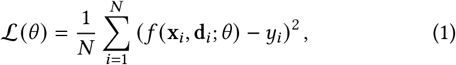

where *N* denotes the number of training instances.

### 2.2 Dataset Preprocessing

Fig. 1 presents the preprocessing pipeline. CDS regions and genomic sequences were obtained from organism-specific reference annotations. For the Auxin datasets, CDS regions were extracted from Ensembl Gene annotations *Arabidopsis_thaliana.TAIR*10.60 [11], and the genomic sequences were paired with the corresponding ribosome density profiles. For each CDS, we construct windows centered at codon position *i* with total window size *w*. Each window spans codon indices [*i* − ⌊*w* / 2⌋, *i* + ⌈*w* / 2⌉ −1], covering both upstream and downstream contexts. Each training instance consists of the nucleotide sequence within the window, the aligned reference density profile, and the codon-level target value at the center position, obtained by aggregating densities over the corresponding codon. Both target and reference density values are log-transformed to reduce right skewness and stabilize variance.

**Figure 1:**
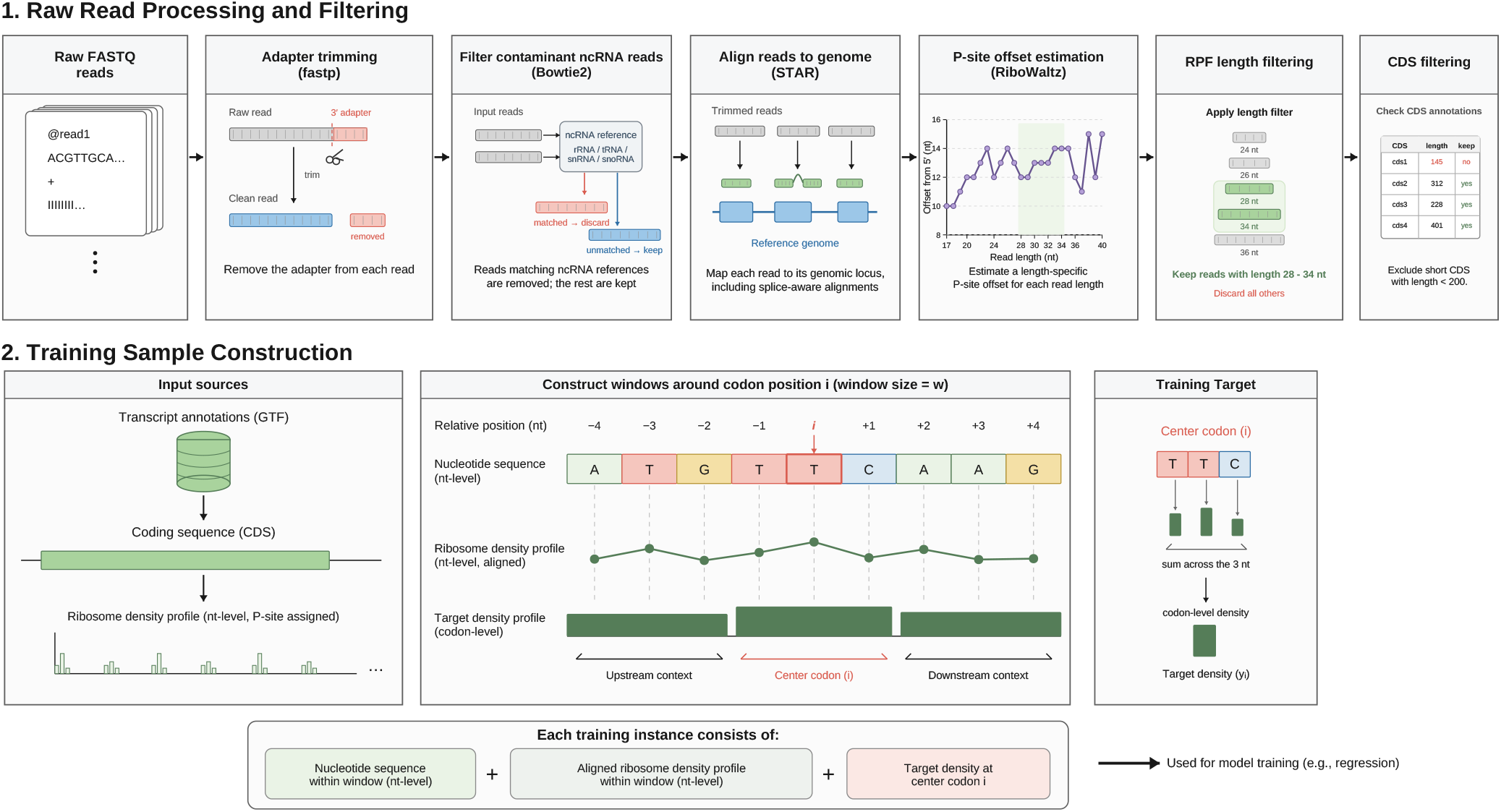
Raw FASTQ reads were processed using a standard pipeline. Adapter trimming was performed with fastp, where reads with more than 25% of bases having quality scores below 25 were removed. Reads mapping to noncoding RNA references were filtered using Bowtie2. The remaining reads were aligned to the reference genome using STAR, followed by P-site offset estimation with RiboWaltz. Read-length filtering was performed separately for each dataset according to its experimental protocol and footprint population. For the Auxin datasets, only ribosome-protected fragments with lengths between 28 and 34 nt were retained. Following [6], we excluded the first and last 10 codons of each Coding Sequence (CDS) to avoid boundary effects. CDS shorter than 200 nt were excluded, and CDS with ribosome coverage below the 75th percentile were filtered to ensure signal quality.

P-site-assigned Ribo-seq data are sparse and highly variable, with many positions having zero or low observed density. To address this, we apply a *log1p* transformation to the target values. Let *d*_*i*_ denote the codon-level ribosome density at position *i*, obtained by aggregating nucleotide-level densities over the corresponding codon. We define the target as:

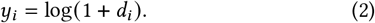

This formulation stabilizes variance and reduces the impact of noise in low-coverage regions. The same transformation is also applied to the reference density profile within each window.

### 2.3 Dual-Stream Input Representation

Fig. 2 illustrates the overall architecture. Each nucleotide *s*_*i*_ is mapped to a *d*-dimensional vector space through an embedding function *f*_seq_ : *X* → ℝ^*d*^. In parallel, the ribosome density values d = (*d*_1_, *d*_2_, …, *d*_*L*_), where *d*_*i*_ ∈ ℝ_≥0_, are projected into the same *d*-dimensional hidden space via a projection function *f*_*d*_ : ℝ_≥0_ → ℝ^*d*^. Both streams are normalized by layer normalization.

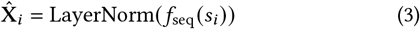

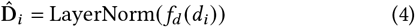

**Figure 2:**
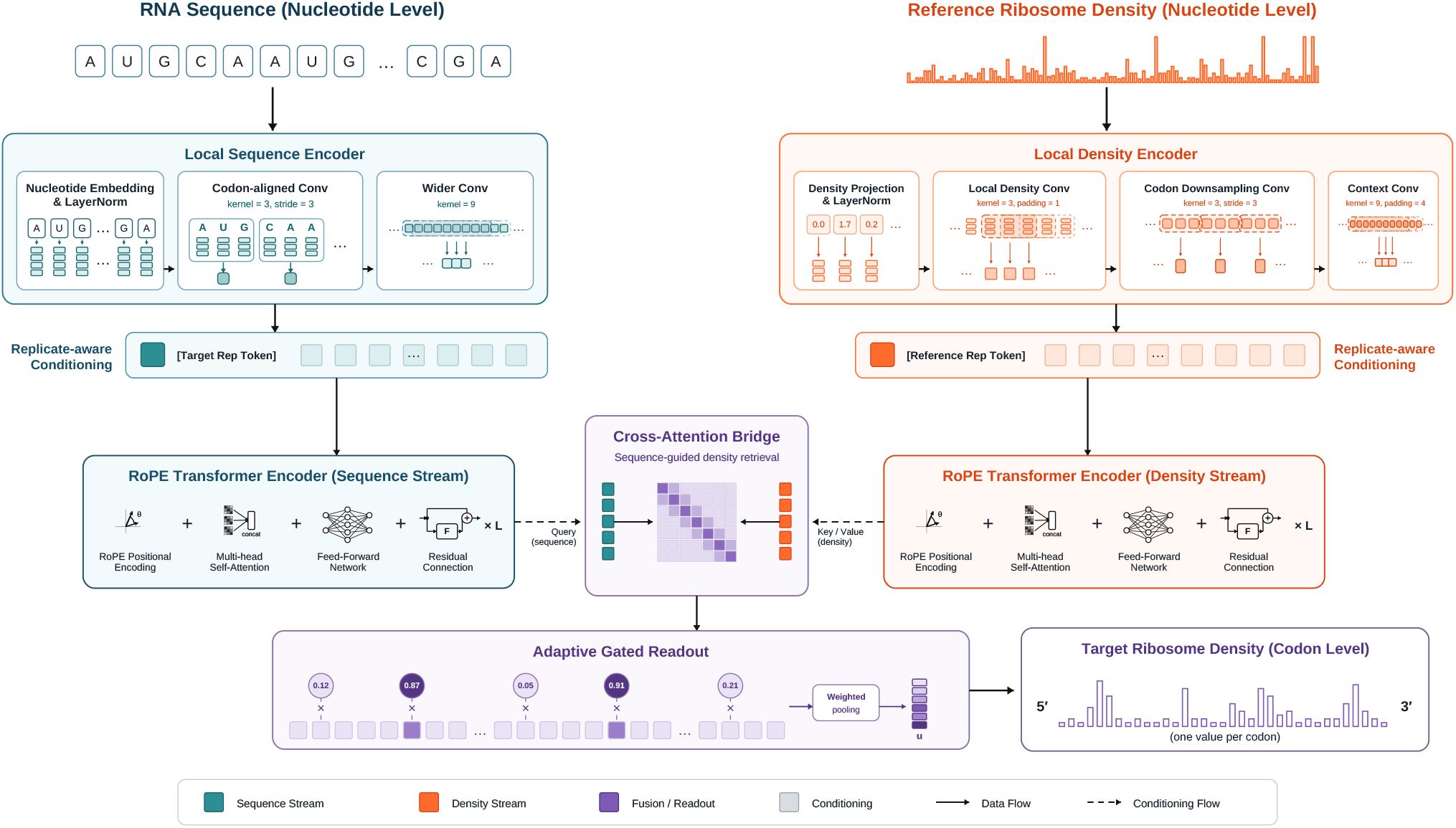
The model takes two aligned input streams at nucleotide resolution, where each nucleotide in the RNA sequence is paired with its corresponding reference ribosome density value. These two modalities are first embedded into a shared hidden space and encoded by branch-specific convolutional blocks to capture short-range patterns while producing codon-resolution representations. To support replicate-specific prediction, we introduce learnable replicate tokens: a target replicate token is prepended to the sequence stream, and a reference replicate token is prepended to the density stream. This allows the model to condition its prediction on replicate identity while preserving separate sequence and density representations. The sequence and density streams are then independently contextualized using RoPE-based Transformer encoder layers. To integrate the two modalities, we apply asymmetric sequence-density cross-attention, where the target-conditioned sequence stream queries the reference-conditioned density stream. The gated readout then computes position-specific weights from the concatenation of sequence-context, density-context, and cross-attended codon representations, while applying the weights to the cross-attended codon representations. The resulting pooled representation is layer-normalized and passed through a linear prediction head to produce the final codon-centered density estimate. Overall, the model consists of three stages: (1) dual-stream representation of sequence and reference density, (2) local and global feature extraction via convolution and RoPE self-attention, and (3) replicate-aware sequence-query cross-modal integration with gated readout for prediction.

#### 2.3.1 Sequence branch

Ribosome profiling signals are sparse and highly variable at nucleotide resolution, making direct modeling prone to noise. In contrast, translation occurs at the codon level, where triplets of nucleotides jointly determine elongation dynamics. As a result, nucleotide-level modeling can be overly fine-grained and may obscure codon-level signals that regulate translation.

To address these challenges, we introduce branch-specific convolutional encoders to extract local patterns from both the sequence and density inputs prior to global feature extraction. Convolution naturally aggregates neighboring nucleotides into short-range representations. Let 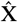 denote the embedded sequence input and 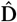 the projected density input.

The sequence branch is processed by a codon-aligned convolutional stack, where a kernel size of 3 and stride of 3 convert the nucleotide sequence into codon-resolution tokens, followed by a wider kernel to capture longer local context:

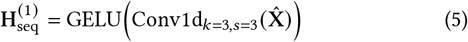

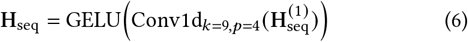

#### 2.3.2 Density branch

Ribosome profiling signals do not correspond to clean codon-aligned measurements and are affected by P-site offset uncertainty, frame ambiguity, and mapping noise, resulting in noisy and imperfect alignment at nucleotide resolution. To accommodate these properties, the density branch first operates at nucleotide resolution to preserve fine-grained variations without prematurely enforcing codon alignment, followed by a local convolution to smooth noise in the ribosome profiling signal. The smoothed signal is then aggregated using a stride-3 convolution, where each step groups neighboring nucleotides into 3-mer (codon-level) units, introducing a codon-level inductive bias and reorganizing the signal into codon-resolution representations. Finally, a wider convolution expands the receptive field to capture local dependencies across multiple codons.

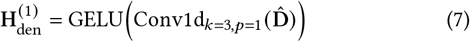

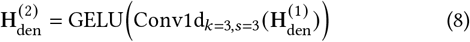

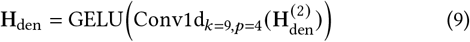

### 2.4 Replicate-Aware Transformer

#### 2.4.1 Replicate-aware conditioning

We introduce replicate embeddings as task-conditioning tokens to explicitly encode replicate context. This is analogous to the use of special tokens and prompts in transformer-based language models [9, 10, 12]. Let *r*_target_ and *r*_ref_ denote the target and reference replicate identifiers, respectively. After local convolution, a target replicate token is prepended to the sequence stream, while a reference replicate token is prepended to the density stream:

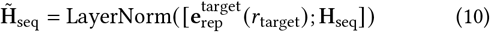

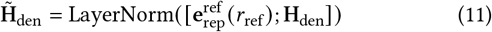

The replicate tokens provide global conditioning signals that are visible to all positions during subsequent self-attention.

#### 2.4.2 Translational context attention

Following RoFormer [13], we adopt rotary positional embeddings (RoPE) to encode positional information directly within the attention. Unlike additive absolute positional embeddings, RoPE makes query-key interactions depend on relative positional offsets without adding absolute positional embeddings. For a generic stream input H = (h_1_, …, h_*L*_), where h_*m*_ ∈ ℝ^*d*^ denotes the representation of the *m*-th token after replicate-aware conditioning, RoPE-based self-attention is independently applied to the sequence and density streams to contextualize intra-stream representations:

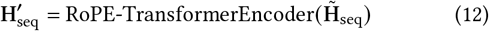

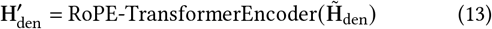

Each encoder layer uses pre-normalized self-attention followed by a feed-forward network with residual connections. For each attention head, positional information is injected into the query and key representations by rotating them according to their positions:

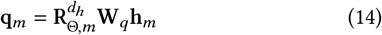

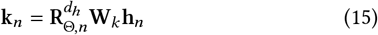

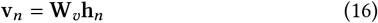

where *d*_*h*_ is the head dimension, and 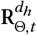 denotes the block-diagonal rotary matrix at position *t*. The inverse frequencies are predefined as 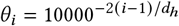. The resulting query-key interaction can be written as

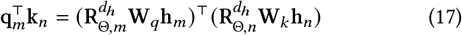

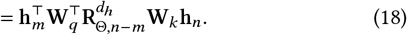

The contextualized representation at position *m* is computed as

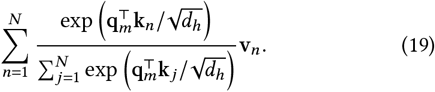

#### 2.4.3 Sequence-Density Cross-Attention

After intra-stream contextualization, the two branches encode complementary aspects of the same window. The sequence branch represents the target-conditioned cis-regulatory context, while the density branch represents reference-conditioned experimental ribosome occupancy. To integrate these modalities, we use asymmetric cross-attention in which the sequence stream queries the density stream:

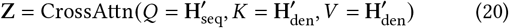

The RNA sequence is shared between the reference and target replicates and defines the codon context at which translation is predicted. The reference density profile provides experimental evidence about ribosome occupancy under the reference condition. By using the sequence stream as the query and the density stream as the key and value, the model learns which reference-density positions are most informative for each sequence-derived translational context. The resulting representation Z incorporates density-derived evidence while preserving the sequence stream as the primary representation for predicting target-replicate ribosome density.

#### 2.4.4 Gated readout

After sequence-density cross-attention, the model obtains a representation that integrates nucleotide context with reference ribosome-density information. Because different positions within the input window may contribute unequally to the center-codon prediction, we use a gated readout network to adaptively weight position-wise representations before readout.

Let s_*i*_, d_*i*_, and c_*i*_ ∈ ℝ^*d*^ denote the sequence-context, density-context, and cross-attended representations at codon position *i*, respectively. The replicate conditioning tokens are excluded from this readout step, so the gate is computed only over codon-level positions. For each codon position, we first concatenate the three representations and normalize them:

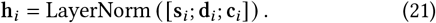

A scalar gate is then computed as

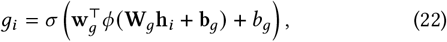

where *ϕ*(·) denotes the GELU activation [14] and *σ*(·) is the sigmoid function. The final readout is obtained by applying the gate to the cross-attended representation and computing a normalized weighted mean:

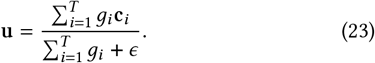

The pooled vector is then layer-normalized and passed to a linear prediction head:

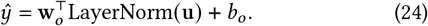

## 3 Experiments

We performed five-fold cross-validation: each fold held one partition as test set, one for validation, and three for training. Mean squared error was used as the training objective, and the model achieving the lowest validation loss was selected and evaluated on the corresponding test set. Models were trained on an NVIDIA RTX 5080 GPU (16 GB) using the AdamW optimizer with a cosine annealing learning rate schedule initialized at 5 × 10^−4^, weight decay of 0.01, and gradient clipping with a maximum norm of 1.0. Training was performed for up to 50 epochs with early stopping based on validation loss using a patience of 5 epochs. All models were implemented in PyTorch 2.11.0 with Python 3.13.

### 3.1 Datasets

The model was evaluated using ribosome profiling datasets from three distinct experimental settings, covering bacterial, yeast, and plant systems. These datasets were selected to assess whether the proposed model can learn relationships between ribosome occupancy profiles across different biological conditions, experimental protocols, and ribosome-associated footprint populations.

First, the GSE119104 dataset [15] was used to evaluate cross-protocol prediction in Escherichia coli. This dataset compares ribosome profiling signals generated using Cm-lysis and high-Mg protocols. The Cm-lysis data were used as the reference input, while the high-Mg data were used as the prediction target. This task was designed to test whether the model can learn systematic differences between ribosome profiling protocols and recover the target-condition ribosome occupancy profile from a related reference profile.

Second, the GSE139036 dataset [16] was used to evaluate prediction between different ribosome footprint populations in Saccharomyces cerevisiae. In this setting, monosome ribosome footprint signals were used as the reference input, and disome footprint signals were used as the prediction target. Because monosome and disome footprints represent distinct ribosome-associated populations, this task evaluates whether information contained in standard monosome ribosome occupancy profiles can be used to predict regions enriched for disome-associated ribosome density.

Third, the Auxin datasets were used to evaluate model performance on plant ribosome profiling data from Arabidopsis thaliana seedlings exposed to a rapid auxin perturbation. In these experiments, seedlings were treated with auxin for 30 seconds, providing a short-timescale condition intended to capture early translational responses before extensive downstream transcriptional or developmental changes occur. The analysis included two biological replicate datasets, Auxin-R1 and Auxin-R2 as well as a combined Auxin-R1/R2 setting. The single-replicate settings were used to evaluate model performance within each replicate, whereas the combined setting was used to test whether jointly modeling multiple biological replicates could improve prediction by capturing shared auxin-associated ribosome occupancy patterns while accounting for replicate-specific variation.

For all datasets, codon-centered sequence windows of 120 nucleotides were used as model input. A dropout rate of 0.1 was applied during training to reduce overfitting. Hyperparameters were selected by grid search over the hidden dimension, number of attention heads, and number of transformer layers. The search space and selected settings are summarized in Table 2.

**Table 1:** Dataset statistics for the experimental datasets. For each gene, we used a canonical isoform policy and did not allow alternative splicing. #CDS candidates denotes CDS regions that passed the minimum length filter and excluded mitochondrial chromosomes. #Representative CDS denotes CDS regions retained after canonical isoform selection. #CDS Final denotes representative CDS regions that generated at least one valid codon-centered window after boundary-codon exclusion and ribosome-density filtering. #Windows denotes the final number of codon-centered model input windows.

| Study | #Ref<br>Replicates | #Target<br>Replicates | #Genes<br>Annotated | #Genes<br>Used | #CDS<br>Candidates | #CDS<br>Retained | #CDS<br>Final | #Windows |
| --- | --- | --- | --- | --- | --- | --- | --- | --- |
| GSE119104 (*E. coli*, Cm-lysis → High-Mg) | 1 | 1 | 8,591 | 1,026 | 4,150 | 4,105 | 1,026 | 269,963 |
| GSE139036 (*S. cerevisiae*, monosome → disome) | 1 | 1 | 6,600 | 1,590 | 6,372 | 6,372 | 1,590 | 626,862 |
| Auxin-R1 | 1 | 1 | 27,655 | 6,737 | 47,434 | 26,942 | 6,737 | 2,351,136 |
| Auxin-R2 | 1 | 1 | 27,655 | 6,737 | 47,434 | 26,942 | 6,737 | 2,357,783 |
| Auxin-R1/R2 | 2 | 2 | 27,655 | 6,737 | 47,434 | 26,942 | 6,737 | 9,404,544 |

**Table 2:** Hyperparameters used in the baselines.

| Hyperparameter | Value |
| --- | --- |
| Hidden dimension ($d$) | [16, 32, 48] |
| Attention heads | [2, 4, 8] |
| Transformer layers | [1, 2, 4] |

### 3.2 Baselines

We compare against five baselines. For RiboMIMO, Riboexp, and Riboformer, we implemented window-level baselines based on the corresponding source code.^2^

- **CNN [17]**: A four-layer 1D convolutional network with 128 filters per layer and no attention mechanism.
- **TF-Enc [7]**: A vanilla transformer encoder that takes the sequence and reference-density streams as inputs, adds positional embeddings, and models the codon-centered window with self-attention.
- **RiboMIMO [4]**: We preserve the original BiGRU feature extractor and dual prediction heads. The regression head predicts the ribosome-density value at the center codon, while the classification head follows the original RiboMIMO fast/slow/pausing density-state definition. We adapt the full-CDS multi-output formulation to the codon-centered window prediction setting.
- **Riboexp [5]**: We preserve the original policy-network idea, where a neural network learns to select codons informative for ribosome-density prediction, but adapt the full-CDS formulation to the codon-centered window prediction setting.
- **Riboformer [6]**: This method applies convolutional encoders to the sequence input and reference-density input independently, extracting local sequence motifs and density-context features. The encoded features are then modeled with self-attention to capture contextual dependencies across the codon-centered window. We reimplemented the source code in PyTorch.
- **RiboRep**. Our proposed model, including local convolutional feature extraction, replicate-aware conditioning, RoPE-based Transformer encoding, sequence-density cross-attention, and gated readout.

### 3.3 Experimental Results

Table 3 shows model performance across datasets. We evaluate performance using a gene-level data-splitting strategy. All windows derived from the same gene are assigned to the same training, validation, or test set. This provides a stricter evaluation because the model must generalize to genes not observed during training.

**Table 3:** Model performance is evaluated using Pearson correlation, Spearman correlation, and MSE.

| Study | Model | Pearson | Spearman | MSE |
| --- | --- | --- | --- | --- |
| Auxin-R1 | CNN | 0.5988 ± 0.0151 | 0.3644 ± 0.0048 | 0.0497 ± 0.0051 |
| Auxin-R1 | TF-Enc | 0.6402 ± 0.0167 | 0.3942 ± 0.0042 | 0.0459 ± 0.0044 |
| Auxin-R1 | Riboexp | <u>0.6645 ± 0.0173</u> | 0.3907 ± 0.0034 | <u>0.0437 ± 0.0037</u> |
| Auxin-R1 | RiboMIMO | 0.2675 ± 0.0180 | 0.2437 ± 0.0043 | 0.0720 ± 0.0091 |
| Auxin-R1 | Riboformer | 0.6530 ± 0.0161 | <u>0.4014 ± 0.0046</u> | 0.0445 ± 0.0043 |
| Auxin-R1 | RiboRep | **0.6887 ± 0.0159** | **0.4074 ± 0.0063** | **0.0427 ± 0.0044** |
| Auxin-R2 | CNN | 0.5872 ± 0.0232 | 0.3346 ± 0.0037 | 0.0518 ± 0.0054 |
| Auxin-R2 | TF-Enc | 0.6031 ± 0.0193 | 0.3521 ± 0.0051 | 0.0504 ± 0.0059 |
| Auxin-R2 | Riboexp | 0.6116 ± 0.0184 | 0.3428 ± 0.0027 | 0.0495 ± 0.0051 |
| Auxin-R2 | RiboMIMO | 0.2529 ± 0.0134 | 0.2236 ± 0.0033 | 0.0740 ± 0.0091 |
| Auxin-R2 | Riboformer | <u>0.6136 ± 0.0259</u> | <u>0.3567 ± 0.0048</u> | <u>0.0492 ± 0.0048</u> |
| Auxin-R2 | RiboRep | **0.6592 ± 0.0283** | **0.3620 ± 0.0047** | **0.0443 ± 0.0035** |
| Auxin-R1/R2 | CNN | 0.5935 ± 0.0217 | 0.3496 ± 0.0038 | 0.0506 ± 0.0046 |
| Auxin-R1/R2 | TF-Enc | 0.6273 ± 0.0232 | 0.3754 ± 0.0037 | 0.0478 ± 0.0060 |
| Auxin-R1/R2 | Riboexp | <u>0.6475 ± 0.0224</u> | 0.3719 ± 0.0171 | <u>0.0452 ± 0.0031</u> |
| Auxin-R1/R2 | RiboMIMO | 0.2415 ± 0.0174 | 0.2268 ± 0.0033 | 0.0738 ± 0.0084 |
| Auxin-R1/R2 | Riboformer | 0.6289 ± 0.0138 | <u>0.3792 ± 0.0050</u> | 0.0473 ± 0.0048 |
| Auxin-R1/R2 | RiboRep | **0.6667 ± 0.0207** | **0.3852 ± 0.0066** | **0.0446 ± 0.0043** |
| GSE119104 | CNN | 0.7339 ± 0.0178 | 0.7481 ± 0.0163 | 1.5468 ± 0.0313 |
| GSE119104 | TF-Enc | 0.8025 ± 0.0136 | 0.8295 ± 0.0133 | 1.1927 ± 0.0265 |
| GSE119104 | Riboexp | <u>0.9230 ± 0.0085</u> | <u>0.8364 ± 0.0120</u> | <u>0.0940 ± 0.0035</u> |
| GSE119104 | Riboformer | <u>0.9196 ± 0.0099</u> | 0.8342 ± 0.0114 | 0.0983 ± 0.0051 |
| GSE119104 | RiboMIMO | 0.6066 ± 0.0225 | 0.4874 ± 0.0365 | 0.4079 ± 0.0372 |
| GSE119104 | RiboRep | **0.9231 ± 0.0086** | **0.8421 ± 0.0093** | **0.0938 ± 0.0035** |
| GSE139036 | CNN | 0.5833 ± 0.0269 | 0.3692 ± 0.0194 | 0.2131 ± 0.0103 |
| GSE139036 | TF-Enc | <u>0.6718 ± 0.0260</u> | <u>0.4728 ± 0.0198</u> | **0.1769 ± 0.0082** |
| GSE139036 | Riboexp | 0.6619 ± 0.0267 | 0.4669 ± 0.0180 | 0.1864 ± 0.0100 |
| GSE139036 | RiboMIMO | 0.5521 ± 0.0327 | 0.4285 ± 0.0187 | 0.2304 ± 0.0116 |
| GSE139036 | Riboformer | **0.6728 ± 0.0244** | **0.4760 ± 0.0154** | <u>0.1787 ± 0.0073</u> |
| GSE139036 | RiboRep | 0.6525 ± 0.0260 | 0.4389 ± 0.0162 | 0.1859 ± 0.0106 |

#### 3.3.1 Auxin datasets

RiboRep shows the best performance across Auxin-R1, Auxin-R2, Auxin-R1/R2 datasets. The model consistently improves prediction correlations over the baselines across both single-replicate and combined-replicate settings. For Auxin-R1, RiboRep achieved the best Pearson correlation. Similar improvements are observed for Auxin-R2 and the combined Auxin-R1/R2 setting.

These gains indicate that the proposed replicate-aware architecture is particularly effective when multiple biological replicates are jointly modeled. The improvement is most pronounced in Pearson correlation, while Spearman improvements are more moderate. This suggests that the model better captures quantitative density variation, whereas rank-level ordering remains challenging. Overall, the results support that replicate-aware conditioning and gated readout improve generalization on plant ribosome profiling data, especially under the stricter gene-level split.

#### 3.3.2 GSE119104 dataset

This experiment was designed to evaluate whether the models can recover protocol-dependent changes in ribosome occupancy patterns from ribosome profiling data generated under Cm-based and High Mg harvesting protocols. Previous studies have reported protocol-dependent ribosome pausing differences between Cm-based and High Mg harvesting procedures [6, 15]. RiboRep achieved the best mean performance across all three evaluation metrics on GSE119104, with the highest Pearson correlation and Spearman correlation and the lowest MSE. However, its Pearson correlation and MSE were nearly tied with those of Riboexp, whereas the improvement in Spearman correlation was more noticeable. These results indicate that RiboRep is competitive with the strongest baseline for cross-protocol ribosome-density prediction and may better preserve the relative ordering of ribosome-occupancy values.

#### 3.3.3 GSE139036

The results on GSE139036 were more mixed. Riboformer achieved the highest Pearson and Spearman correlations, while TF-Enc obtained the lowest MSE, although the two models showed very similar overall performance, with only marginal differences across all three metrics. RiboRep remained competitive but did not outperform the strongest baselines on this dataset. These results suggest that predicting disome ribosome-density profiles from monosome profiles may require additional biological information or dataset-specific modeling beyond the current framework.

### 3.4 Ablation Study

To evaluate the effectiveness of each proposed component in RiboRep, we compare the full model against four ablated variants. The analysis focuses on four aspects: local feature extraction, replicate-aware conditioning, relative positional encoding, and gated readout. Table 4 reports the experimental results of the ablated variants across different datasets.

**Table 4:** Ablation study of the RiboRep components evaluated using Pearson correlation, Spearman correlation, and MSE.

| Study | Model | Pearson | Spearman | MSE |
| --- | --- | --- | --- | --- |
| Auxin-R1 | RiboRep | **0.6887 ± 0.0159** | **0.4074 ± 0.0063** | **0.0427 ± 0.0044** |
| Auxin-R1 | RiboRep (w/o Conv) | 0.5820 ± 0.0249 | 0.3536 ± 0.0139 | 0.0509 ± 0.0035 |
| Auxin-R1 | RiboRep (w/o RoPE) | 0.5328 ± 0.0031 | 0.3299 ± 0.0025 | 0.0487 ± 0.0017 |
| Auxin-R1 | RiboRep (w/o Gate) | <u>0.6589 ± 0.0472</u> | <u>0.3924 ± 0.0218</u> | <u>0.0436 ± 0.0042</u> |
| Auxin-R2 | RiboRep | **0.6592 ± 0.0283** | **0.3620 ± 0.0047** | **0.0443 ± 0.0035** |
| Auxin-R2 | RiboRep (w/o Conv) | 0.5598 ± 0.0247 | 0.3222 ± 0.0059 | 0.0538 ± 0.0038 |
| Auxin-R2 | RiboRep (w/o RoPE) | 0.5385 ± 0.0133 | 0.3156 ± 0.0060 | 0.0538 ± 0.0001 |
| Auxin-R2 | RiboRep (w/o Gate) | <u>0.6412 ± 0.0405</u> | <u>0.3524 ± 0.0130</u> | <u>0.0460 ± 0.0043</u> |
| Auxin-R1/R2 | RiboRep | **0.6667 ± 0.0207** | **0.3852 ± 0.0066** | 0.0446 ± 0.0043 |
| Auxin-R1/R2 | RiboRep (w/o Conv) | 0.5835 ± 0.0342 | 0.3504 ± 0.0168 | 0.0510 ± 0.0030 |
| Auxin-R1/R2 | RiboRep (w/o RoPE) | 0.5522 ± 0.0343 | 0.3280 ± 0.0092 | 0.0539 ± 0.0035 |
| Auxin-R1/R2 | RiboRep (w/o Gate) | <u>0.6570 ± 0.0243</u> | <u>0.3843 ± 0.0028</u> | <u>0.0445 ± 0.0049</u> |
| Auxin-R1/R2 | RiboRep (w/o Rep) | 0.6526 ± 0.0229 | 0.3795 ± 0.0089 | **0.0445 ± 0.0029** |
| GSE119104 | RiboRep | **0.9231 ± 0.0086** | **0.8421 ± 0.0093** | **0.0938 ± 0.0035** |
| GSE119104 | RiboRep (w/o Conv) | <u>0.9062 ± 0.0072</u> | <u>0.8112 ± 0.0149</u> | <u>0.1117 ± 0.0040</u> |
| GSE119104 | RiboRep (w/o RoPE) | 0.8512 ± 0.0115 | 0.7264 ± 0.0202 | 0.1721 ± 0.0081 |
| GSE119104 | RiboRep (w/o Gate) | 0.9004 ± 0.0060 | 0.7952 ± 0.0199 | 0.1216 ± 0.0161 |
| GSE139036 | RiboRep | **0.6525 ± 0.0260** | <u>0.4389 ± 0.0162</u> | **0.1859 ± 0.0106** |
| GSE139036 | RiboRep (w/o Conv) | 0.6354 ± 0.0286 | **0.4409 ± 0.0255** | 0.1956 ± 0.0115 |
| GSE139036 | RiboRep (w/o RoPE) | 0.6361 ± 0.0239 | 0.4374 ± 0.0219 | 0.2069 ± 0.0137 |
| GSE139036 | RiboRep (w/o Gate) | <u>0.6473 ± 0.0268</u> | 0.4371 ± 0.0277 | <u>0.1888 ± 0.0103</u> |

- **RiboRep (w/o Conv)**. This variant removes the local convolutional encoder and relies directly on the transformer to model nucleotide-level dependencies.
- **RiboRep (w/o RoPE)**. This variant removes rotary positional embeddings from the transformer encoder. It tests the importance of relative positional information for modeling codon-context dependencies.
- **RiboRep (w/o Gate)**. This variant removes the gated read-out module and replaces it with a mean-pooling readout. It evaluates whether adaptive feature selection over codon-level representations improves prediction performance.
- **RiboRep (w/o Rep)**. This variant removes replicate identity conditioning. It evaluates whether explicitly encoding reference and target replicate identities improves replicate-specific density prediction.

#### 3.4.1 Effectiveness of Local Feature Extraction

This ablation evaluated the importance of local feature extraction before global attention. Removing the convolutional encoder degraded Pearson correlation and MSE across all datasets and reduced Spearman correlation in most settings, with the exception of a slight increase on GSE139036. Overall, these results indicate that local convolutional feature extraction contributes substantially to predictive performance. This observation is biologically plausible, as translation is fundamentally a codon-level process. The convolutional encoder introduces an explicit locality bias before global self attention is applied. In addition, ribosome profiling signals are often sparse and noisy at nucleotide resolution. The convolutional encoder aggregates local nucleotide context and ribosome density signals to capture local density patterns and reduce sensitivity to position level noise. This may also help preserve local reading frame structure and three nucleotide periodicity in ribosome profiling signals.

#### 3.4.2 Effectiveness of Replicate-Aware Conditioning

This ablation evaluated the importance of replicate-aware conditioning. We report results only on Auxin-R1/R2, the only dataset in our benchmark containing multiple replicates. When multiple reference and target replicates are available, we use all possible reference-target pairings during both training and evaluation. In Auxin-R1/R2, this results in four combinations: Control-R1 to Treatment-R1, Control-R1 to Treatment-R2, Control-R2 to Treatment-R1, and Control-R2 to Treatment-R2.

To evaluate the contribution of replicate-aware conditioning, we remove the special reference and target replicate tokens, thereby withholding explicit information about the replicate identities of the two profiles. As shown in Table 4, removing replicate-aware conditioning resulted in modest decreases in the Pearson and Spearman correlations, whereas the MSE remained comparable to that of the full model. These results suggest that replicate-aware conditioning may help the model better capture the overall variation and relative ordering of ribosome densities.

#### 3.4.3 Effectiveness of Relative Positional Encoding

This ablation was designed to investigate the importance of positional information in ribosome density prediction. The substantial performance drop compared to other variants suggests that positional information is crucial for modeling ribosome density profiles. Without positional encoding, the transformer becomes permutation-invariant and cannot distinguish whether a motif or signal occurs near or far from the P-site.

#### 3.4.4 Effectiveness of Gated Readout

This ablation was designed to assess the contribution of adaptive readout weighting. We hypothesized that different positions within the window contribute unequally to the center-codon prediction, and that uniform mean pooling may dilute informative local signals. This design is conceptually related to Riboexp [5], which uses a policy network to identify codons that are informative. The gated readout assigns position-specific importance weights before aggregation. The consistent improvement over the mean-pooling variant suggests that adaptively emphasizing informative positions while suppressing less relevant contextual representations benefits ribosome-density prediction.

## 4 Discussion

RiboRep builds on the reference-guided prediction setting introduced by Riboformer [6]. Accordingly, its primary distinction does not lie in the reference-guided prediction task itself, but in its nucleotide-level sequence representation and cross-attention-based integration of sequence and ribosome-density information.

Beyond the experiments evaluated in this study, RiboRep could potentially be extended to broader downstream applications demonstrated by Riboformer, including cross-condition ribosome-profile prediction, prediction and interpretation of ribosome-collision profiles, and correction of protocol-dependent experimental biases [6]. These applications would, however, require task-specific training and independent validation.

For example, sequence patterns associated with ribosome pausing could be investigated by computing Sequence Impact Scores (SIS) through in silico sequence perturbation, following the approach used in Riboformer. Alternatively, input-attribution methods such as Integrated Gradients [18] could be applied to identify nucleotide or codon positions that contribute strongly to predicted ribosome pausing. High-attribution subsequences could then be aggregated and analyzed to identify candidate sequence motifs associated with the predicted pausing patterns.

One potential direction for future work is to compare attribution patterns across replicate combinations to identify nucleotide or codon positions that consistently contribute to predicted ribosome pausing. Such replicate-consistent signals may help distinguish robust sequence determinants from replicate-dependent variation and facilitate the identification of reproducible candidate motifs.

## 5 Conclusion

In this work, we presented RiboRep, a replicate-aware cross-modal transformer framework for codon-resolved ribosome density prediction. Unlike prior approaches that primarily operate on codon-level representations, RiboRep preserves nucleotide-resolution sequence and ribosome occupancy signals during early feature extraction and integrates reference ribosome-density information through asymmetric sequence-density cross-attention. We further introduced replicate-aware conditioning to model biological and technical variation across replicates.

Across bacterial, yeast, and plant ribosome profiling datasets, RiboRep achieved competitive or improved performance compared with existing baselines. The most consistent improvements were observed on the Auxin datasets and the E. coli protocol-comparison task, supporting the effectiveness of replicate-aware and sequence-density integrated modeling. Performance on the S. cerevisiae monosome-to-disome prediction was more mixed, suggesting that more complex ribosome-associated states may require additional biological context beyond local sequence and occupancy patterns. Overall, RiboRep provides a biologically informed and flexible framework for ribosome density prediction across organisms, experimental protocols, and replicate settings. Its improved modeling of translational dynamics may support downstream applications such as identifying translation regulatory elements, characterizing ribosome pausing, and guiding synthetic biology and sequence engineering design. More broadly, RiboRep establishes a foundation for translation-aware molecular digital twins, where codon-resolution ribosome occupancy landscapes can be computationally reconstructed and simulated under diverse biological and perturbation conditions. Such digital twin frameworks may enable in silico modeling of translational state transitions, prediction of perturbation-induced protein synthesis responses, and integration of translational regulation into multiscale cellular-state simulation. Future work will incorporate additional biological signals, including codon optimality, tRNA abundance, mRNA secondary structure, and RNA-binding protein interactions, to further improve robustness, interpretability, biological fidelity, and cross-condition generalization.

## 6 Acknowledgment

This research is partially supported by the National Academies of Sciences, Engineering, and Medicine, grant number SCON-10001538 to Z.Y. ChatGPT was used to assist in generating the figures based on finalized drafts prepared by the authors. All figures were subsequently revised by the authors. ChatGPT was also used to assist with refactoring the released code, which was subsequently reviewed and validated by the authors.

## Footnotes

1 indicates the corresponding author

2 RiboMIMO: https://github.com/tiantz17/RiboMIMO; Riboexp: https://github.com/Liuxg16/Riboexp; Riboformer: https://github.com/lingxusb/Riboformer.

